## Supplementary figures and images for "Constructing the first comorbidity networks in companion dogs in the Dog Aging Project"

### Supplemental Figure 1

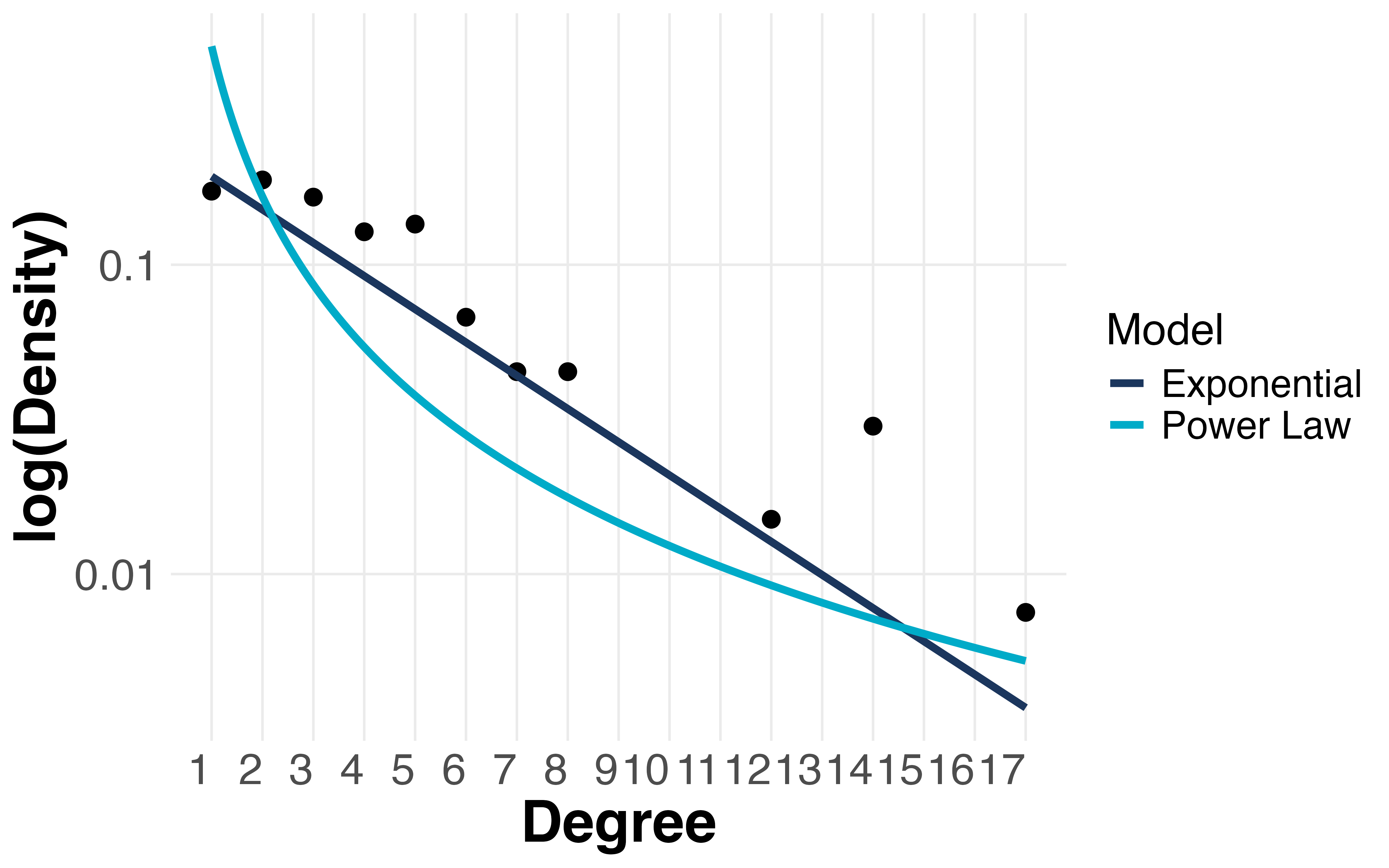
